# Human papillomavirus type 16 sub-lineages and integration in cancer

**DOI:** 10.1101/2020.06.25.162958

**Authors:** Robert Jackson, Alejandro Ortigas-Vásquez, Ingeborg Zehbe

**Author notes:** **Corresponding author:** Robert Jackson, PhD. **Affiliation history:** Robert Jackson was affiliated with Lakehead University and the Thunder Bay Regional Health Research Institute at the time the study was performed and is currently affiliated with the University of Arizona.

## Abstract

Our lab has been intrigued by the fact that viral genomes often take on the role of mobile elements to perpetuate their existence in a complex organism’s genome. Multiple DNA viruses such as Epstein-Barr virus, hepatitis B virus, and human papillomavirus (HPV) can invade their host genome, as “genomic parasites”. We have been investigating HPV type 16 (HPV16), which is a prominent human tumour virus. In our recent *in vitro* work using 3D organoids, a common variant of HPV16’s coding region elicited early integration into the host genome compared to the HPV16 prototype sequence. Next-generation sequencing (NGS) data confirmed a transcriptomic profile of increased proliferation and chromosomal instability—both hallmarks of cancer. Epidemiologically, this variant is associated with a high cervical cancer incidence. To take inquiries a step further, we investigated variant-specific integration across HPV16-related cancers using NGS data from population-derived clinical samples in The Cancer Genome Atlas (TCGA)-curated database. Data were analyzed for HPV16 positivity, sub-lineage, and viral-host integration using a bioinformatic pipeline of open-source tools, including HPVDetector. Here, we report the analysis of 120 cervical cancer cases comprising HPV16 positive and negative samples as well as their different sub-lineage and integration status. The integration signature between variant and prototype did not differ quantitatively but qualitatively: that of the variant being related to hypoxia/energetics (Warburg effect) and that of the prototype being much more varied to include host immune abrogation and cancer pathways activation. We conclude by discussing challenges and future directions for expanding these analyses.

## INTRODUCTION

The co-evolutionary interplay between pathogens and their hosts involves a plethora of intriguing biological phenomena, including genomic integration of pathogen DNA into host cells. While cellular integration of viral genomic material is a means of propagation for families such as *Retroviridae* (*e*.*g*., human immunodeficiency virus, HIV), it can also occur during persistent infections with DNA viruses and is often associated with virally induced cancers. One such family of common DNA tumour viruses, *Papillomaviridae*, is responsible for causing ∼5% of all human cancers worldwide [Ghittoni *et al*., 2015]. There are more than 600 types of papillomaviruses (PVs) discovered as of July 2021 based on the PaVE reference genome database [PaVE: https://pave.niaid.nih.gov, Van Doorslaer *et al*., 2013; 2017a]. The number of PV types, including the non-human PVs (currently 226) is expected to continue rising due to high-throughput genomics and sampling [Van Doorslaer and Dillner, 2019]. The majority of PVs currently identified are the human papillomaviruses (HPVs) [Van Doorslaer *et al*., 2017b] and these types (448 to date) differ in their propensity for inducing cancer as well as their epithelial tropism, with “high-risk” types being the primary cause of malignancies: predominantly via sexually-transmitted persistent HPV type 16 (HPV16) infection of anogenital and oropharyngeal mucosa [Ndiaye *et al*., 2014]. Although this and other HPV genomes are considered “small”, only ∼7.9 kb containing nine open reading frames (ORFs) and a variety of alternatively spliced and multi-cistronic variant transcripts [Graham & Faizo, 2017], it encodes potent oncoproteins such as E6 and E7 which drive host cells to acquire the hallmarks of cancer [Hanahan and Weinberg, 2000; 2011; Mesri *et al*., 2014]. The combined E6 and E7 actions disable protective apoptotic mechanisms and promote host cell proliferation as outlined by Hoppe-Seyler and colleagues [2018]—a phenomenon that could potentially be exploited therapeutically at an RNA [Togtema *et al*., 2018] or protein-level [Togtema *et al*., 2019]. Of particular relevance for this study, the HPV-augmented environment can permit chromosomal instability (an enabling characteristic among the hallmarks of cancer) and eventual integration of HPV16 sequences into human DNA via double-strand break (DSB) repair events [Winder *et al*., 2007], yielding unique integration signatures found in next-generation sequencing (NGS) data from human carcinomas [Holmes *et al*., 2016].

HPV16 sub-lineages (variants) via epidemiological and lab-based studies have been reported to infer differing cancer risk [reviewed by Mirabello *et al*., 2018 and references within; Clifford *et al*., 2019]. HPV16 variant designations were originally based on their geographical region of origin and have since been updated to lineage (1.0 to 10.0% whole viral genome sequence difference) and sub-lineage (0.5 to 1.0% difference) nomenclature. The European “prototype” sub-lineage (EP; now A1 according to new nomenclature) was the first HPV16 genome published [Seedorf *et al*., 1985]. The remaining sub-lineages include two additional European groups (E; now A2 and A3), one Asian (As; now A4), eight African (Af-1a, -1b and −2; now B1-4 and C1-4), two North-American (NA; now D1 and D4) and two Asian-American (AA-1 and −2; now D3 and D2, respectively) [Burk *et al*., 2013; Mirabello *et al*., 2018]. We have put our efforts toward comparing the EP and AA sub-lineage variants, which differ by only three amino acid changes at residues 14 (Q to H), 78 (H to Y) and 83 (L to V) in the major transforming protein E6 [Richard *et al*., 2010; Niccoli *et al*., 2012; Jackson *et al*., 2014; 2016; Togtema *et al*., 2015; Cuninghame *et al*., 2017]. Epidemiological studies revealed that the AA sub-lineage infers a higher risk for dysplasia and an earlier onset of invasive tumours than EP [Xi *et al*., 1997; Berumen *et al*., 2001]. Our functional assays showed that AAE6 has a greater transforming, migratory, and invasive potential than EPE6 when the respective E6 gene was retrovirally transduced into primary human keratinocytes in long-term *in vitro* immortalization studies [Richard *et al*., 2010; Niccoli *et al*., 2012; Togtema *et al*., 2015]. These observations may be due, at least in part, to an AAE6-mediated altered metabolic phenotype reminiscent of the Warburg effect [Richard *et al*., 2010; Cuninghame *et al*., 2017]. Consistent with the hypothesis that AAE6 is more oncogenic than EPE6, our recent experimental work of early HPV16 carcinogenesis revealed that AAE6 in the context of the full-length viral genome is more prone to host genome integration than is EPE6 [Jackson *et al*., 2014; 2016]. At that time, one study mentioned HPV16 sub-lineage variants and integration but did not find a difference (*P* = 0.28, two-tailed Fisher’s exact test) between EPE6 (3 episomal and 20 integrated cases) and the E-T350G variant (6 episomal and 16 integrated cases, responsible for one of the residue changes also found in AAE6: L83V) [Xu *et al*., 2013]. Notably, however, the lone tumour sample that contained the AA variant was in integrated form.

Based on our assumption that the propensity for HPV16 integration may vary between sub-lineages, we set out to do a search of clinical samples with available genomic and transcriptomic data reported in databases. Two landmark publications of The Cancer Genome Atlas (TCGA) reported on cervical squamous cell carcinoma and endocervical adenocarcinoma (CESC) [TCGA Research Network, 2017] and head and neck squamous cell carcinoma (HNSC) [TCGA Research Network, 2015]. More recently, Cantalupo *et al*. [2018] reported further analysis of additional TCGA data, including additional cancer sets that have become available in recent years. They also used a clearly defined, rules-based approach to filtering and detection: first extracting high-quality non-human reads followed by removal of non-relevant viral families (bacteriophage, plant viruses, insect viruses) and artifacts, with 10 or more high-quality alignments required for virus detection. Prior to the primary TCGA data analysis papers, there was a publication about DNA viruses in TCGA RNA-Seq data across a variety of human cancers [Khoury *et al*., 2013]. An additional literature search found several more related papers: HPV and HNSC comprehensive analysis [Castellsagué *et al*., 2016] and an HPV16-focused follow-up TCGA-HNSC analysis [Nulton *et al*., 2017]. In the latter, three different groupings of integration status were reported: episomal, integrated, and human-viral episomal hybrids, challenging integration data by Parfenov *et al*. [2014]. The Parfenov *et al*. study disclosed HPV typing and sub-lineage analyses of the 35 HPV+ HNSC cases with 29 containing HPV16: one of which was an AA variant. Interestingly, integration with 16 breakpoints in the human genome (*i*.*e*., ∼ 4-fold more compared to EP) was shown. In the Nulton *et al*. study, for the same case, a new phenomenon of human-viral hybrids in the host genomic DNA subsequently found in episomal form was reported [Nulton *et al*., 2017]. Recently, Zapatka *et al*. [2020] further analyzed TCGA data to determine viral associations with cancer, including the impact of integrations on host copy number variations. Most recently, Lou *et al*. [2020] report that integration rates indeed differ between sub-lineages in cervical cancers. In a Guatemalan population with a 24%, 31%, and 45% prevalence of D2, D3, and A1 sub-lineages, respectively, they found that D2 had a higher rate of integration than D3 (76% vs 44%), coupled with an earlier age of cancer diagnosis. Interestingly, they also found that A1 had an equally high rate of integration (78%) as D2, which is in contrast with our *in vitro* data (at least, within this specific population) [Jackson et al., 2016]. It is noteworthy that A1 is phylogenetically distant from D2/D3, with ∼2% genome difference (>150 single-nucleotide polymorphisms, SNPs, and insertions/deletions, indels), whereas D2 and D3 differ from each other by only ∼0.5% (<40 SNPs and indels, and hence it remains open what the biological meaning for these findings may be).

Our overall objective was to screen TCGA for all HPV16-containing samples, verify their sub-lineage genotypes, then detect and characterize instances of viral integration along with a complete picture of its impact on the host genome and identify potential contributing factors in the surrounding molecular landscape. The most pertinent research question addressed here is whether there are specific integration mechanisms associated with different HPV16 sub-lineage genotypes: is the AA variant (D2 and D3) more frequently associated with integration than the EP variant (A1) and/or are there unique integration patterns with potential functional relevance? Such findings would resonate with our previous variant-specific integration work [Jackson *et al*., 2014; 2016] and provide evidence for a novel mechanism in differing variant tumourigenesis. Our approach of HPV16 sub-lineage genotyping and integration detection, along with an attempt to reconcile and verify existing data, has not been previously addressed for TCGA data. We analyzed TCGA cervical cancer cases (*n* = 120), including DNA-and RNA-level data for each (*n* = 240 total samples). We did not find significantly more or less integration events between AA and EP samples but differences in the integration pattern. AA—in a singular manner—seems to adapt host cells to a hypoxia/energetics (Warburg effect)-associated phenotype whereas EP alters host cells to various hallmarks of cancer including host immune surveillance and cancer pathway signaling. A variety of bioinformatics pipelines exist for sub-lineage genotyping and integration detection. These are however susceptible to biological (*e*.*g*., low copies and incomplete viral coverage) and technical (*e*.*g*., inadequate read depth) hurdles, which are important considerations for future analyses.

## RESULTS & DISCUSSION

### HPV16 D2/D3 integration data are limited for cervical and head and neck cancers

When seeking to test the relationship between HPV16 sub-lineage and integration propensity we started by compiling a sample summary of previously published and available data from TCGA [**Table 1**]. Next, we performed a preliminary sub-lineage and integration analysis for available CESC sample data [**Figure 1**] as it was stratified into sub-lineage for the A lineage, whereas HNSC data were not further stratified between A1, A2, and A3. Overall, while the proportion of integrated cases was 1.23x higher in D2/D3-containing CESC samples (9/10, 90.0% vs 51/70, 72.9% for A1), the sample size was very low rendering this initial analysis underpowered and the finding as not statistically significant (*P* = 0.242, Χ^2^ test).

**Table 1.**
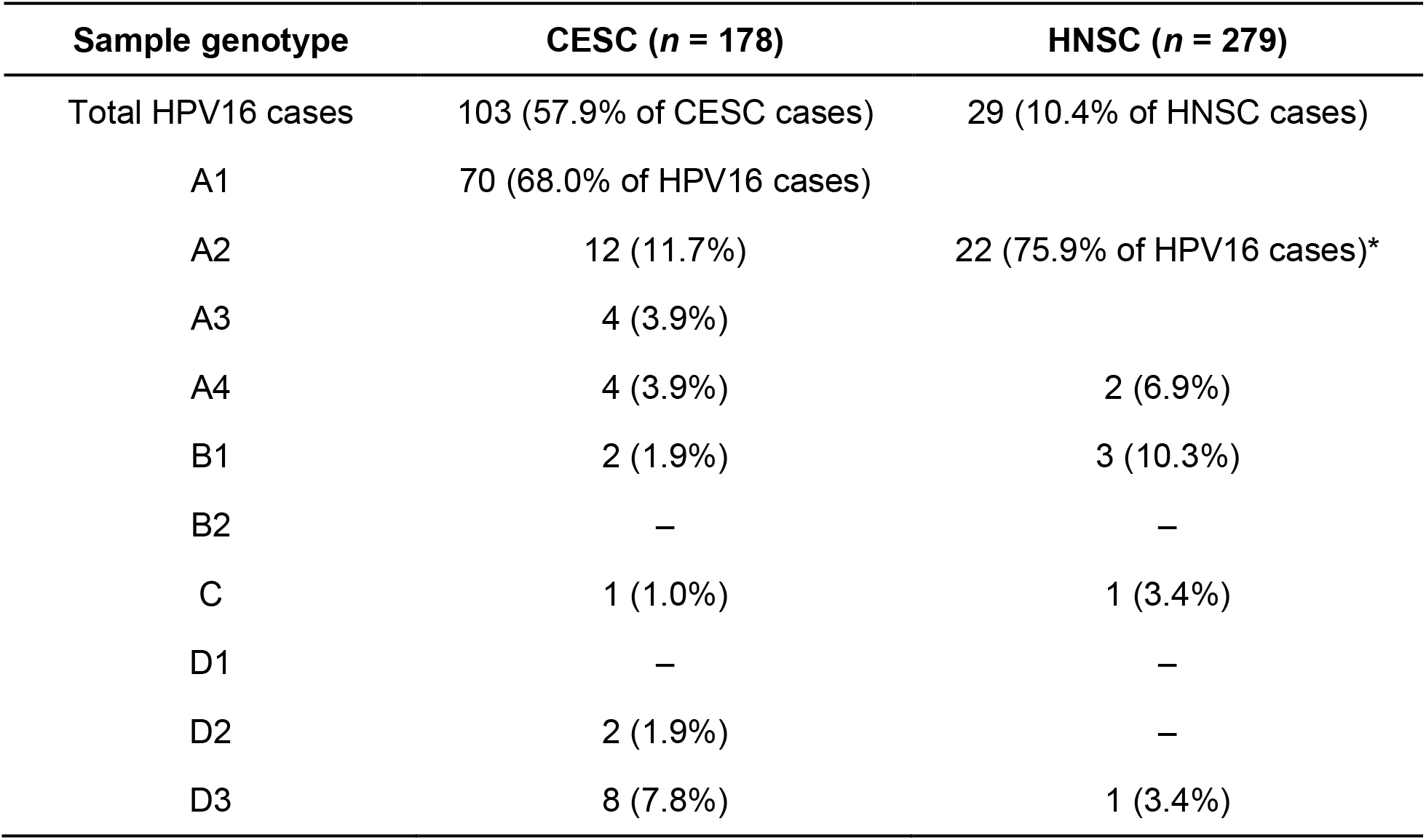
Sample summary breakdown for each TCGA case set. Sample genotypes for cancer cases from the Cervical Squamous Cell Carcinoma and Endocervical Adenocarcinoma (CESC) as well as Head and Neck Squamous Cell Carcinoma (HNSC) datasets. CESC HPV16 data are based on TCGA Research Network [2017] (using the sub-lineages identified at the time, which were only 10 of the current 16). *HNSC HPV16 data are based on Parfenov *et al*. [2014], where the A sub-lineage was grouped as “EUR” genotypes without specifying A1, A2 or A3.

**Figure 1.**
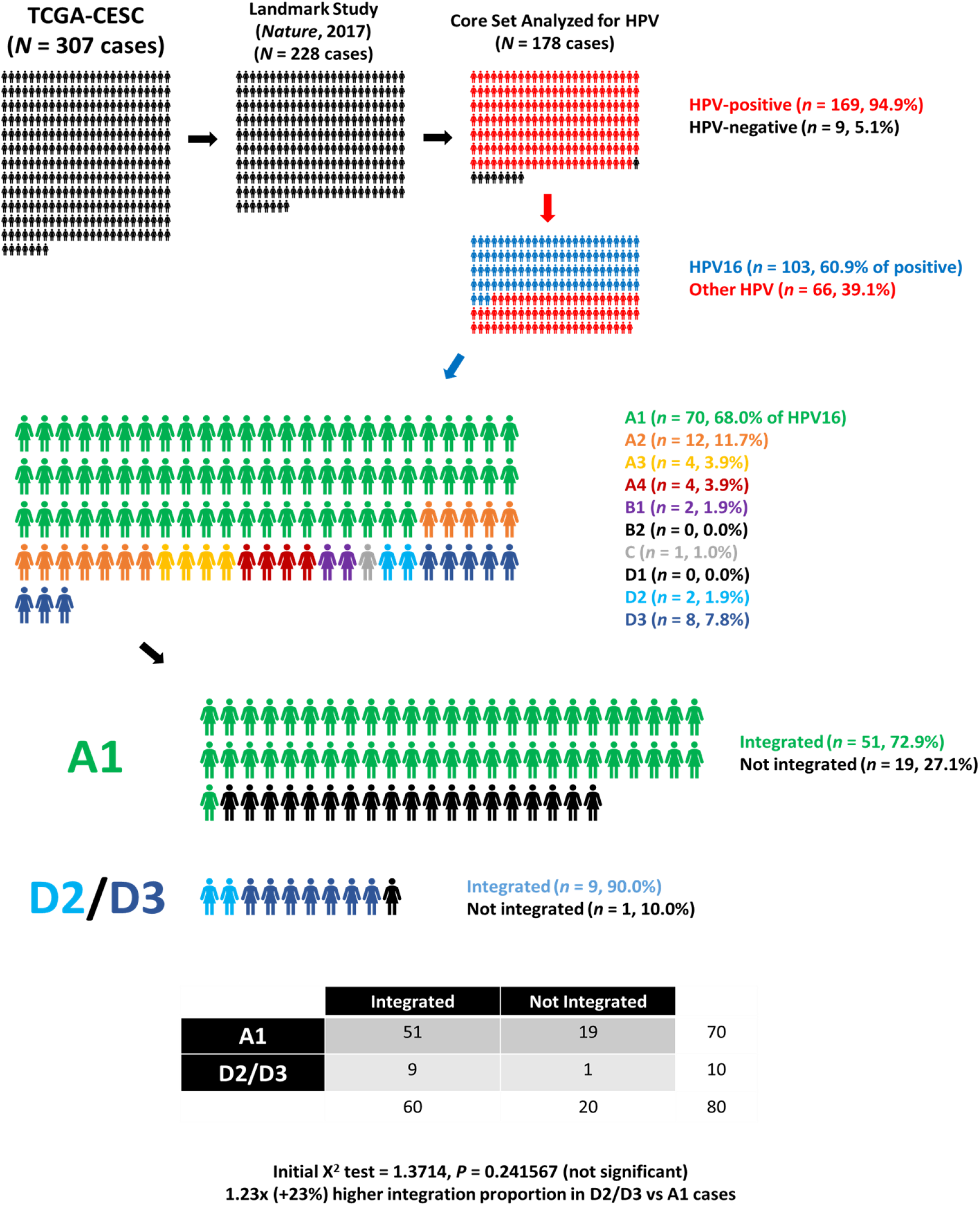
Preliminary analysis of CESC samples. Previously available HPV16 sub-lineage and integration data for the CESC dataset, published by the TCGA Research Network [2017], were analyzed to determine whether there was a higher proportion of integrated samples corresponding to D2/D3 (which have viral genomes that differ from each other by ∼0.5%, but have identical E6 amino-acid sequences) vs the A1 sub-lineage (which differs ∼2% from D2/D3 viral genomes). While these sub-lineage classifications were based on 10 reference genomes, the set has since been expanded to 16 sub-lineages, as described in Mirabello *et al*. [2018].

### The untapped potential of TCGA

To expand upon these preliminary results, we were using our own HPV16 detection and sub-lineage analyses to verify sub-lineage calls by prior studies, resolve discrepancies, and further analyze the “unknown” samples. From the National Cancer Institute’s Genomic Data Commons (GDC) Data Portal, there are currently 835 relevant CESC (*n* = 307) and HNSC cases (*n* = 528) available for analysis, including 2,568 binary alignment map (BAM) files (27.84 TB total, an average of 33.34 GB/case, including all whole-exome sequencing (WXS) and RNA-Seq sequencing read files and excluding miRNA-Seq files as they are not primarily relevant for HPV16 genotyping). There are 307 CESC cases (924 BAM files, 10.44 TB) and 528 HNSC cases (1,644 BAM files, 17.41 TB), with both sets including samples from primary tumours, metastases, normal blood, and normal solid tissues. It appears the TCGA legacy data portal also contains whole-genome sequencing (WGS) files for these cancer sets. Based on the TCGA landmark studies [TCGA Research Network, 2015; 2017], we expect the majority of CESC samples are HPV positive, whereas a smaller proportion of HNSC samples are HPV positive. While HPV16 is expected to be the predominant type, multi-type infections and co-infections with other non-commensal microorganisms are also possible. As with most nucleic-acid based techniques, cross-contamination between samples is a potential source of confusion, especially for adjacent cases where sequence libraries could be contaminated [Cantalupo *et al*., 2015].

The number of CESC and HNSC samples that were previously marked as being positive for HPV16 differ between studies, likely due to differences in analysis pipelines (*e*.*g*., data types analyzed, thresholding, and filtering for contamination) [TCGA Research Network, 2017; Cantalupo *et al*., 2018]. While both RNA-Seq and WXS were used for integration analysis in the landmark CESC study, it seems that the authors mostly if not totally relied on RNA-Seq [TCGA Research Network, 2017]. Moreover, for integration status analysis, an in-house tool was used, which cannot be directly compared with other published and available tools such as ViFi [Nguyen *et al*., 2018], HPVDetector [Chandrani *et al*., 2015] or VirTect [Xia *et al*., 2019] (and others detailed in the **METHODS** section). This poses major challenges to a current, valid analysis of sub-lineage association with integration based on these previous publications.

### HPV16 positivity quantification reveals discrepancy between RNA-Seq and WXS samples

We analyzed both RNA-Seq and WXS CESC samples independently to detect the presence of HPV16 in a subset of 120 cases (240 samples, **Figure 2**) representing all 103 available HPV16 positive cases identified in the landmark study as well as 17 negatives to determine how robust our methods are [**Figure 3A**]. As expected, control sample SiHa was a low-positive (5.51 VRPM) and CaSki was a high-positive (502.66 VRPM). TCGA RNA-Seq samples had a range of 0.00 to 1,182.78 VRPM, whereas WXS samples ranged from 0.00 to 109,994.41 VRPM. In total, 85/120 (70.8%) of the cases we analyzed were HPV16 positive based on RNA-Seq and used for subsequent sub-lineage and integration analysis. However, for the same cases, DNA and RNA level HPV16 quantification were often inconsistent, with only 67/120 (55.8%) cases yielding consistent HPV16 quantification (negative, low-positive, or high-positive).

**Figure 2.**
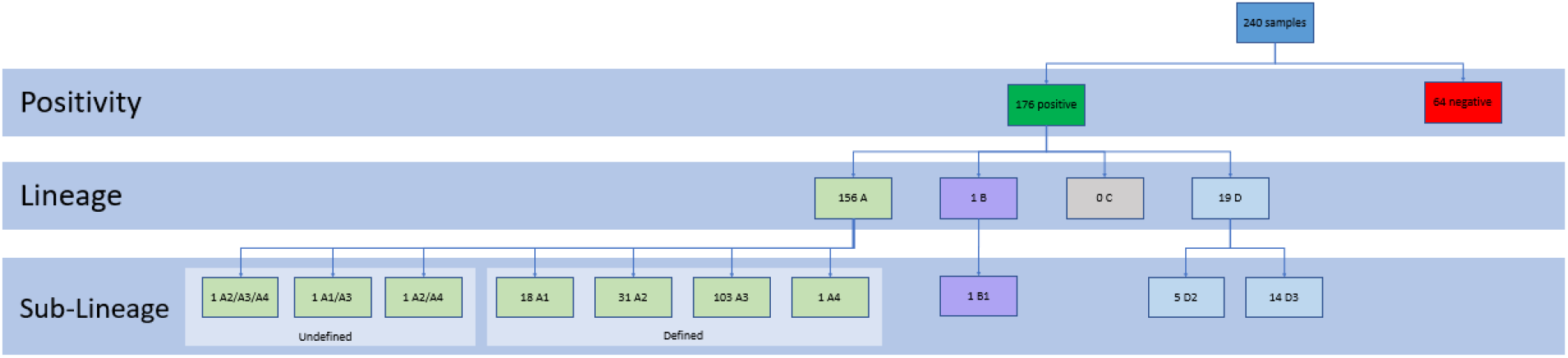
Flowchart summary of all 240 samples analyzed. The 240 samples analyzed were broken down based on positivity (based on RPM), lineage and sub-lineage (based on reference genome with most mapped reads). Samples mapping equally to more than one reference genome were grouped under “Undefined”.

**Figure 3.**
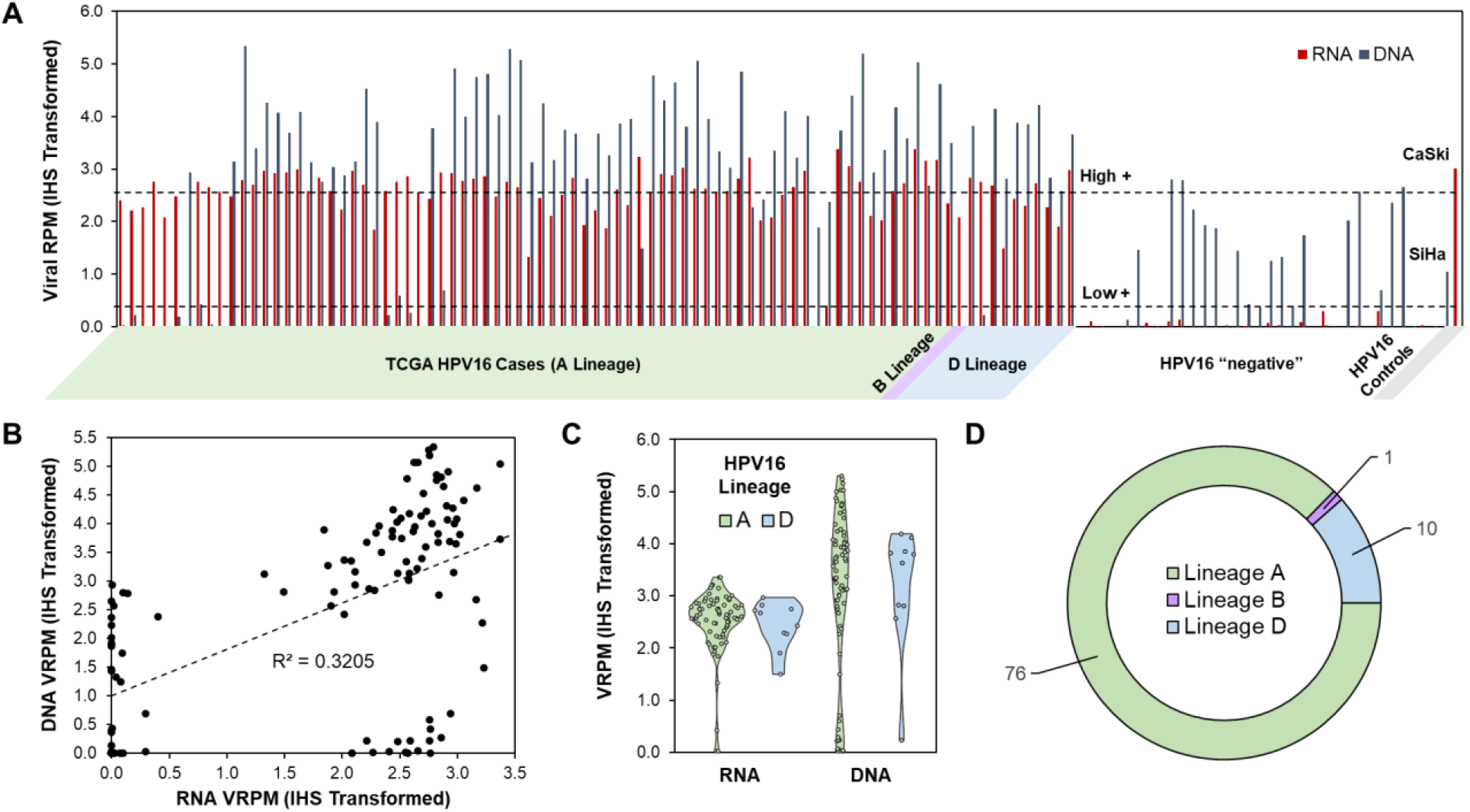
HPV16 positivity in RNA-Seq and WXS TCGA samples. **(A)** A total of 240 samples were analyzed to determine whether they were positive for HPV16, with 120 corresponding to RNA-Seq and 120 corresponding to WXS samples. Samples with less than 0.99 viral reads per million (RPM) (marked by the lowest dashed line) were classified as negative. Positive samples were then subdivided based on strength of positivity, with RPM values below 99 (marked by the highest dashed line) indicating low positivity and those above 99 indicating high positivity. TCGA case IDs are colour-coded based on their CESC landmark study lineage determination (A = green, B = purple, and D = blue). Cervical cancer control cell lines, SiHa and CaSki (shaded grey), were included as low- and high-positive controls, respectively. Values are inverse hyperbolic sine (IHS) transformed. **(B)** Correlation plot of RNA vs DNA-level HPV16 VRPM. **(C)** Sina plot [Sidiropoulos *et al*., 2018] of VRPM for HPV16 lineage A vs D. **(D)** HPV16 lineage pie chart, for the 87 HPV16 positive (RNA-level) cases analyzed.

To resolve these discrepancies, we compared our data for both RNA-Seq and WXS samples to data from the landmark CESC study [TCGA Research Network, 2017], summarized in [**Figure 4**]. We found that positivity results for RNA-Seq samples were the most consistent with previous findings, as all cases that had been reported as negative for HPV16 were also found negative in our analysis at the RNA level, but not necessarily at the DNA level, where we found 7 WXS samples above the low-positive threshold (2 of which are even above the high-positive threshold). Interestingly, 9/10 (90.0%) of the D lineage samples were positive at both the DNA and RNA level, whereas only 62/94 (66.0%) of the A lineage samples analyzed were positive at both DNA and RNA levels. These discrepant findings could be due to biological or technical phenomena, such as insufficient read depth, too few viral reads (small amount of viral DNA in a subset of cells, relative to host DNA), or no/weak transcription of integrated viral sequences (in the cases where RNA-Seq signal is lacking). These issues could be further compounded by computational artefacts and potential contaminating viral sequences, which makes it essential (yet challenging) to develop tools that successfully distinguish contaminating from authentic viral sequences [Cantalupo *et al*., 2018].

**Figure 4.**
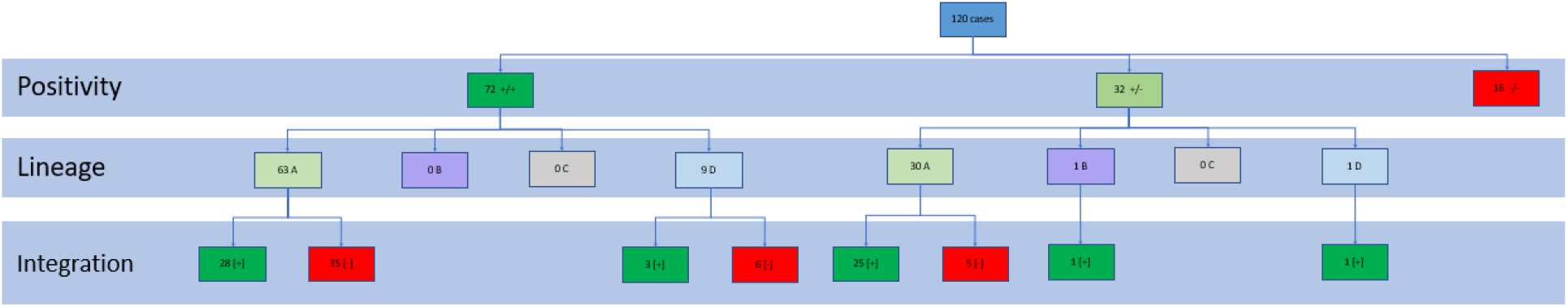
Flowchart summary of all 120 cases analyzed. The 120 cases analyzed were broken down based on positivity [depending on whether both samples were positive (+/+), one was positive and the other negative (+/-), or whether both were negative (-/-)], lineage [defined by both corresponding samples giving a matching lineage call] and integration [based on number of repeated integration reads].

Overall, for this set of 120 cases there was a weak positive correlation between DNA and RNA HPV16 levels (R^2^ = 0.3205) [**Figure 3B**]. Lineage-specific HPV16 levels were compared for DNA and RNA, with both similar between A and D lineages [**Figure 3C**]. The majority of HPV16 positivity was attributable to the A lineage [**Figure 3D**].

### HPV16 lineage, but not sub-lineage, was consistent between RNA-Seq and WXS

Our genotyping results were highly consistency among RNA-Seq and WXS samples at the lineage level, with all sample pairs with matching positivity calls also possessing matching lineage calls. However, of these cases, only 7 matched at the sub-lineage level (1 match for D2 and 6 matches for D3). No exact matches were found at the sub-lineage level for samples of the A or B lineages. This could be due to the high degree of sequence similarity between members of the A lineage as opposed to those belonging to the D lineage, making the sub-lineage determination dependent on a higher degree of difference between references. In the case of WXS sample TCGA-IR-A3LA, our genotyping algorithm was not capable of distinguishing between sub-lineages A2, A3 and A4, indicating equal identity for all three sequences based on mapping statistics. The final lineage distribution of the 72 cases positive for HPV16 can be seen in **Figure 4**. For our controls, SiHa and CaSki were both within A lineage, with SiHa identified as A1/A3 and CaSki as sub-lineage A2. Overall, our results are consistent with others reporting that sub-lineage identification is challenging, with difficulty in confidently assigning a single sub-lineage among highly similar members of the same lineage [Dawson *et al*., 2019].

An important consideration is that accurate sub-lineage identification can be negatively impacted by integration itself, due to truncation of viral genomic regions which may decrease the amount of genomic space available for alignment. Also, it is possible that some integration events may render the viral genome transcriptionally inactive, but still tumourigenic due to insertional mutagenesis (*e*.*g*., disrupting a tumour suppressor). In this theoretical example, DNA-level high-throughput sequencing (*e*.*g*., WGS and WXS) would be required to resolve the integration site, but low sequencing depth and a low abundance of virus relative to host DNA could yield false-negatives for viral positivity, sub-lineage identification, and integration. This is a limitation with TCGA datasets. Viral sequence capture enrichment could address this concern, so long as probes are designed to target across the entire viral genome and flanking host regions.

### The proportion of HPV16 integrated reads does not differ between lineages

Initial raw output indicated the presence of viral integration sites in all samples that had been found positive for HPV16 in earlier steps of the pipeline, suggesting a higher than anticipated false-positive rate. While it is possible these could represent a low proportion of truly integrated reads, we decided that by considering the sequencing depth of each sample we could obtain a more reliable metric in the form of percentage of integrated reads [**Figure 5A**]. Samples were then filtered based on this value and compared to findings from the landmark CESC study [TCGA Research Network, 2017]. Once again, results from RNA-Seq samples were the most consistent with previous reports, indicating presence of integrated HPV16 [**Figure 5A**]. In this case, the proportion of integrated reads was higher in RNA-Seq vs WXS samples. The HPV16 positive cervical carcinoma controls resulted in positive integration for SiHa, while CaSki was considered negative with the current methodology. Clearly, CaSki’s tandemly repeated full genomes pose a problem for detection and represent a false-negative, highlighting the importance of complimentary techniques and a careful biological interpretation and analysis. Integrated reads (%) in HPV16 lineages A vs D were compared and indicate no difference at the RNA or DNA level **[Figure 5B]**, yet a high amount of variability among the HPV16 positive samples.

**Figure 5.**
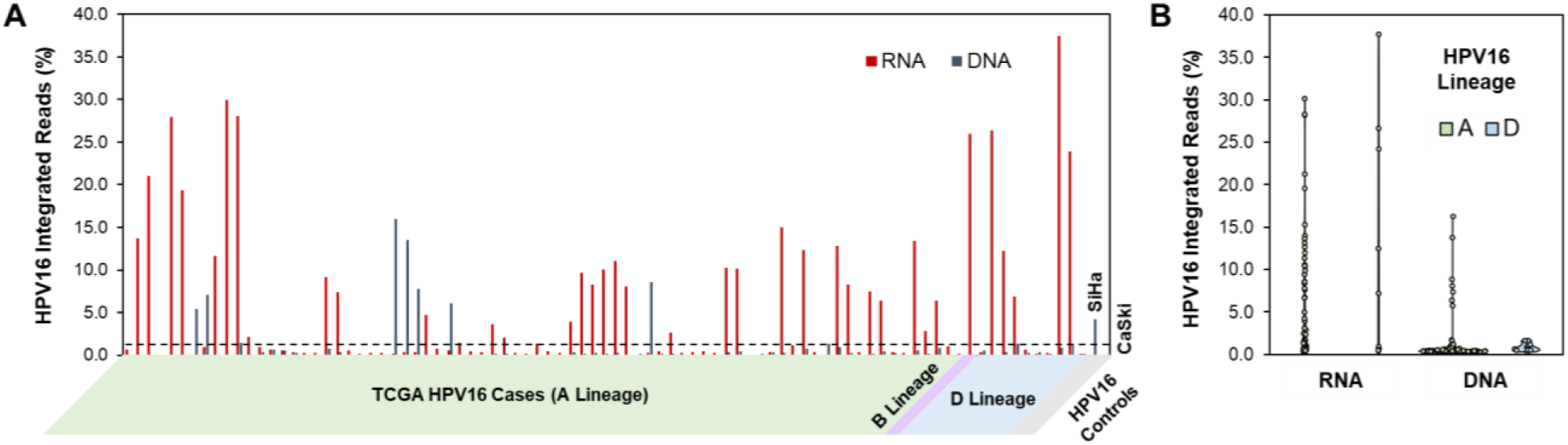
Integrated HPV16 reads in RNA-Seq and WXS samples. **(A)** For each of the HPV16 positive samples, the total number of integrated reads reported by HPVDetector were converted to a percentage of the total number of viral reads. Samples with 1.14% or more integrated reads (marked by the dashed line) were classified as positive for HPV16 integration. Samples below this threshold were classified as predominately containing the episomal form of HPV16. SiHa and CaSki controls were also included, as they are both known to have integrated HPV16. **(B)** Proportion of integrated reads for RNA-Seq and WXS samples comparing HPV16 lineages A and D as a Sina plot [Sidiropoulos *et al*., 2018].

### Integrated genes and known associations with carcinogenesis

Qualitative analysis criteria are based on 63 A lineage and 9 D lineage cases with at least 10 reads in either DNA or RNA-Seq or both [**Figure 4, 6**]. For the A lineages, 28 cases showed corresponding RNA and DNA samples, and for the D lineages, 3/4 cases corresponded, albeit in the 4th case corresponding DNA was found but only with 3 hits. For the A lineages the 31 affected genes were *TAOK3, SUSD1, MIR3134, IKZF3, PLCD3, MAP3K14, IGFBP6, PALLD, VPS13D, IL9R, NFIA, MPRIP, UBA6, UTP11L, DYRK1A, CEACAM5, RXRA, MIPOL1, FLNB, CTSE, FAM220A, ATXN10, RNF166, GRIK4, RAD51B, FHAD1, C6orf203, SIPA1L3, MACROD2, PGAP3, EPHA3*. There are close to 143 genes with more than 10 hits in the DNA-Seq samples with either no or less than 10 RNA-Seq hits, *e*.*g*., *TP73, DLG, MYC* and *RAD51B*. TP73 is particularly interesting as it comes up twice with very high (5342) and low reads (19), respectively. There are 47 genes with more than 10 hits in the RNA-Seq samples. Generally, the remaining DNA-Seq only samples contain less than 100 hits, and with the exception of *MYC*, which shows a corresponding 3 hits in the RNA-Seq, show either no corresponding RNA-Seq data or just 1 hit. It is noteworthy, that the other cases are negative even for such low amounts of hits. In contrast, samples with 10 or more hits in the RNA-Seq often show a corresponding 1 hit of DNA-Seq not only in that particular sample but in several other samples as well. We have not found any real hotspots but earlier-reported *GRIK4* and *RNF166* come up multiple times. For the D lineages, the affected genes were *CUL2, SLC31A1, MRPS28* and *LEPREL1*. While the signature for the A lineages centres around several functions like immune system [*RNF166, CEACAM5, MPRIP, CTSE, IL9R, FAM220A, PALLD, IKZF3*], proto-oncogenes [*PGAP3*], DNA-related processes [*MACROD2, IGFBP6, FHAD1, RXRA, SIPA1L3, UTP11L, NFIA*], cellular processes [*UBA6, FLNB, VPS13D*], signaling [*EPHA3, GRIK4, TAOK3*], and mitochondria [*DYRK1A, C6orf203*], the signature of the D lineages is related to hypoxia/energetics (the Warburg effect). While this may be related to the fewer samples compared to the A lineages, each affected gene was related to this energy axis. This is particularly interesting because it closely relates to earlier findings in our group [Richard *et al*., 2010, Niccoli *et al*., 2012, Cuninghame *et al*., 2017, Dayer *et al*., 2020].

**Figure 6.**
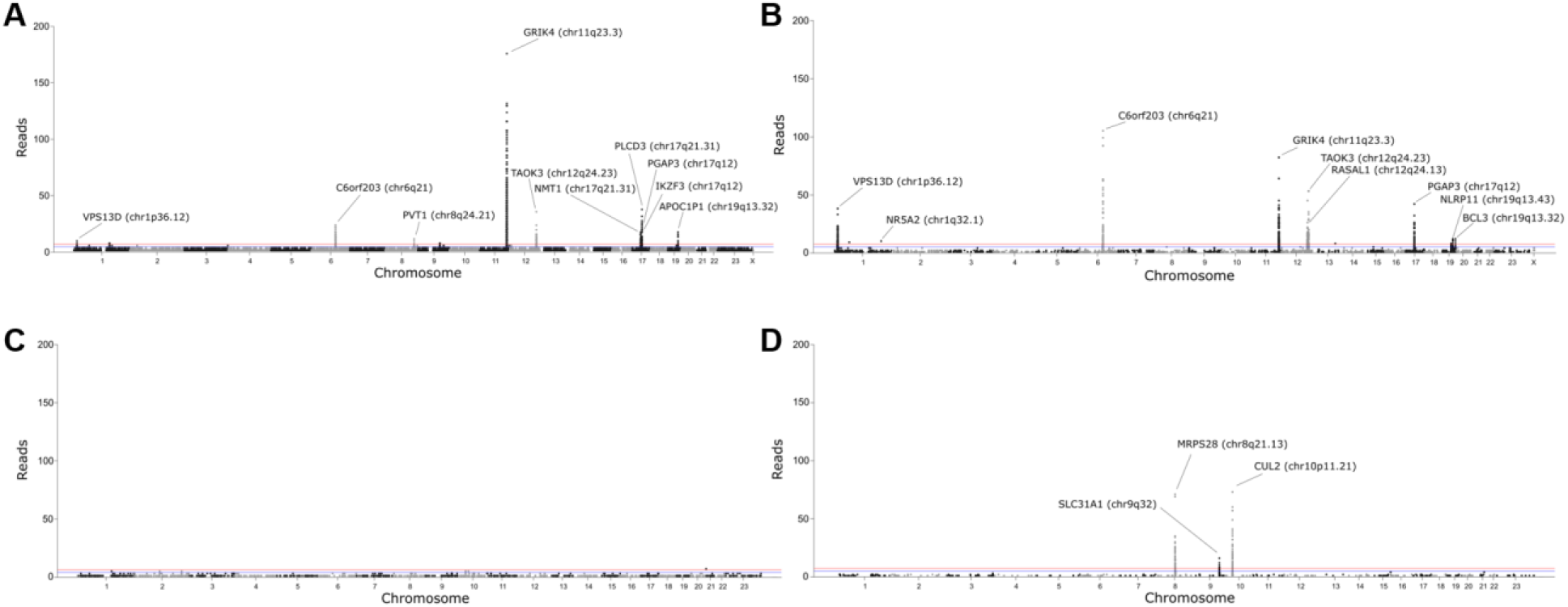
Number of repeated integrated reads per human chromosomal location. Manhattan plots for both A1 and D2/D3 variants based on HPVDetector analyzed RNA-Seq and WXS data indicating the number of repeated integrated reads and the corresponding chromosome number, chromosomal region, and name of the affected gene on the human genome: **(A)** A1 RNA-Seq **(B)** A1 WXS **(C)** D2/D3 RNA-Seq **(D)** D2/D3 WXS. Horizontal lines represent cut-offs: blue = 6 reads; red = 9 reads.

#### HPV16 A1 sub-lineage

Our search with glutamate receptor, ionotropic, kainate 4 encoded by *GRIK4* only retrieved a pharmacogenetic study [Yeon *et al*., 2015] nevertheless associating this neurotransmitter (altered in schizophrenia and depression) in a gene ontology pathway analysis with Wnt/beta-catenin signaling implicated in cancer. Serine/threonine-protein kinase encoded by *TAOK3* was found to be associated with Hippo signaling and a cancer stem cell phenotype [Plouffe *et al*., 2016, Bian *et al*., 2019]. Exons can undergo non-linear reverse splicing to form circ(ular)RNA. Such function has been elucidated in a breast cancer study for Post-GPI attachment to proteins factor 3 encoded by *PGAP3* where overexpressed circPGAP3 inhibited miR-330-3p and upregulated MYC leading to tumour progression and metastases [He *et al*., 2020]. Other studies have also reported a deregulation of PGAP3 in pancreatic [Zhong *et al*., 2020] and breast cancer [Pan *et al*., 2019] without detailing biological consequences but concluding to use PGAP3 respectively as tumour marker or to improve diagnostic accuracy. Mitochondrial transcription rescue factor 1 encoded by *MTRES1* alias *C6orf203* binds highly structured RNA *in vitro* and interacts with the mitochondrial ribosomal large subunit in human cells. Because the loss of C6orf203 leads to reduced mitochondrial translation and consequently, to respiratory inability it was reasoned that C6orf203 is required for efficient mitochondrial translation [Gopalakrishna *et al*., 2019]. Interestingly, C6orf203 showed the highest number of hits in RNA-Seq, which together with DYRK1 could impact A1’s adaption to energetics potentially favouring oxidative phosphorylation (rather than aerobic glycolysis).

#### HPV16 D2/3 sub-lineages

With only 5 breakpoints in the DNA and 4 in the RNA, this sub-lineage showed remarkably fewer hits than the A1 sub-lineage. Interestingly, both CUL2 and MRPS28 proteins are associated with closely-related hallmarks of cancer: hypoxia and deregulated energetics where the altered metabolism “prepares” malignant cells to the hypoxic tumour environment. MRPS28, normally implicated in mitochondrial metabolism, was reported to be modulated in thyroid tumours [Jacques *et al*., 2013] connecting this finding to the Warburg theory of cancer described in 1926 by Otto Heinrich Warburg: “Stoffwechsel der Tumoren” [Brand, 2010]. Warburg argued that rather than fully respiring in the presence of adequate oxygen cancer cells instead resort to aerobic glycolysis fermenting sugar to lactate (rather than pyruvate) [Vander Heiden, 2009]. A newly sparked interest in the *Warburg effect* in the past decade can be gleaned from close to 3,000 publications on the PubMed server alone. Interestingly, MRPS28 is also associated with adaption to hypoxia [Wang *et al*., 2019]. Another finding in this respect was HPV16 D2/D3 integration into the *CUL2* gene [Faull *et al*., 2019, Fouad *et al*., 2019], which interestingly codes for a E3 ubiquitin ligase that targets pRb for degradation [Lee and Zhou, 2010]. While hypoxia-inducing factor 1-alpha (HIF1A) is also targeted by the CUL2 protein, this only leads to its degradation during *normoxia* but not during *hypoxia* where this transcription factor instead accumulates in the cell [Lee and Zhou, 2010 and references therein]. This underlines our previous observations based on lab data that HIF1A is significantly increased in keratinocytes transduced with D2/D3 E6 compared to A1 E6 in the hypoxic environment [Cuninghame *et al*., 2017]. Notably, tumour hypoxia is one of the most challenging aspects of cancer treatment [Cuninghame *et al*., 2014 and references therein]. In the past 20 years, the importance of copper in carcinogenesis (*e*.*g*., solute carrier family 31 member 1, SLC31A1 alias CTR1) has been gaining attention [Barresi *et al*., 2016]. Copper homeostasis is tightly coupled to GSH redox regulation in the mitochondria [Baker *et al*., 2017]. Metal transporting solute carriers mediate control over the availability of endogenous metal ions (*e*.*g*., copper, iron and zinc) and may have key roles in regulating tumour angiogenesis, cell proliferation, epithelial-to-mesenchymal transition (EMT) and aberrant MAPK and STAT3 signal transduction in cancer [Jong *et al*., 2014]. Indeed, copper was demonstrated to increase tumour growth in mice as an outcome of elevated copper levels in drinking water and the switch to aerobic glycolysis (Warburg effect) may be due in part to insufficient copper supply in the tumour microenvironement [Ishida *et al*., 2013]. Similarly, disintegrin and metalloproteinase domain-containing protein 23 is a non-catalytic protein encoded by *ADAM23* with potential roles in tumour progression related to EMT and metastasis [Zmetakova *et al*., 2019].

Overall, it is tempting to speculate that keratinocytes infected with the HPV16 D2/3 sub-lineage adapt to a hypoxia- and Warburg effect-associated phenotype reminiscent of Warburg’s initial notion that cancer is a mitochondrial metabolic disease. Intriguingly, this underlines our experimental findings using cell culture models [Richard *et al*., 2010, Cuninghame *et al*., 2017]. Moreover, patients with cervical cancer containing D2/D3 sub-lineage HPV16 are 7.7 years younger than patients infected with the A1 sub-lineage [Berumen *et al*., 2001] suggesting a more aggressive tumour evolution for D2/D3. The A1 sub-lineage integration signature on the other hand shows variable traits of carcinogenesis related to cancer pathway signaling, and most importantly the potential to tap into oxidative phosphorylation rather than aerobic glycolysis for its energy supply as evidenced by the high expression level of C6orf203. Notably, Warburg’s notion may not be complete since normal proliferating cells with intact mitochondria frequently use aerobic glycolysis as energetics [Vander Heiden, 2009], and it is likely that mixed phenotypes, *e*.*g*., with partially intact mitochondria exist in malignant tumours [Jia *et al*., 2018].

### Present challenges and future directions

While we have addressed relevant literature, acquired access to TCGA sequencing data, designed and setup a working algorithm, and in CESC samples carried out analyses for the HPV16 genotype, sub-lineages, and integration, continued efforts are required to complete our work. Even with TCGA datasets, a major limitation in answering these questions is the small number of D2/D3 compared to A1 samples, and it may be necessary to extend these analyses beyond the well-curated TCGA database, but more broadly into other databases (*e*.*g*., the Sequence Read Archive, SRA). While additional samples could possibly be found and incorporated into our analyses, these data may lack sufficient metadata to be easily searchable and screened (*e*.*g*., experimental vs clinical data). Since the needs of a specific HPV-based analysis may not have been considered *a priori* when data were collected for those samples, there may be significant variability in data formats, read lengths, depths, and quality. Specifically, whether sequence capture or enrichment for HPV sequences was performed would greatly affect the probability of detecting HPV sequences as previously and independently reported by several groups [Holmes *et al*., 2016; Jackson *et al*., 2016]. Sequencing errors could further confound variant identification, requiring careful mitigation [Stewart *et al*., 2018]. Overall, while these challenges exist, potentially relevant datasets deserve analysis and could be useful contributions toward our research question.

Additional work is also required to determine the factors involved in a HPV16 sub-lineage-specific genomic instability and integration risk, including the role of defective DNA damage repair pathways [Seiwert *et al*., 2015; Ratnaparkhe *et al*., 2018]. It is worthwhile to explore relationships between E6 splice variants and genomic stability [Olmedo-Nieva *et al*., 2018]. Researchers continue to explore novel mechanisms surrounding HPV16 integration, such as “super-enhancer-like elements” [Warburton *et al*., 2018] and interruption of tumour suppressor genes [Zhao *et al*., 2016]. Future research could continue exploring these relationships, such as determining the role of non-coding genome elements in genomic instability (*e*.*g*., lncRNAs) [Munschauer *et al*., 2018], the similarities of viral-human integration with mobile elements and human fusion genes [Imielinski and Ladanyi, 2018], as well as evolutionary perspectives [Petrie *et al*., 2018]. Finally, it remains to be seen how clinically-relevant HPV16 sub-lineage genotyping could be, and whether this could lead to improved care and outcome for patients. One strategy could be to compute individual risk scores based on the co-factors affecting the risk of disease progression [Bastarache *et al*., 2018]. New screening techniques could also be useful, such as urine sampling [Van Keer *et al*., 2018] followed by high-throughput sequencing and HPV genotyping, as well as assessing methylation signatures [Cook *et al*., 2018].

Full characterization of the genomic landscape surrounding detected HPV16 integration sites within human chromosomes is required to comprehensively address our research question, *quantitatively* and *qualitatively*. Having a more qualitative approach, by analyzing the affected genes and potential functional consequences of the HPV integration process, makes us conclude that an investigation beyond quantitative statistical analyses needs to be considered. Using both approaches for biological interpretation are important for avoiding computational reproducibility issues: the “alchemy problem” [Rahimi, 2018]. Further characterization of these data includes analyzing viral and host nucleotide positions, sequence overlap/microhomologies, functional annotations of viral and host features, such as structure and gene proximity (inside/outside, exon/intron), promoters, enhancers, transcription factor binding sites, repeat elements, and proximity to fragile sites. “Virtual ChIP-seq” [Karimzadeh and Hoffman, 2018] could be a useful tool to predict nearby transcription factor binding sites (*e*.*g*., CTCF [Doolittle-Hall *et al*., 2015; Paris *et al*., 2015]). As described previously [Jackson *et al*., 2016], the region surrounding a viral integration site can be scanned for repeat elements, which may be prone to rearrangements, using the UCSC human genome browser RepeatMasker track. Another important aspect for qualitative assessment and interpretation will be to visualize and compare the A1 vs D2/D3 cases; example visualizations exist in the literature [Akagi *et al*., 2014; Tang *et al*., 2013; Holmes *et al*., 2016; Jackson *et al*., 2016; Warburton *et al*., 2018; Lagström *et al*., 2019].

## CONCLUDING REMARKS

From a basic science perspective, big data analysis may aid in unraveling the mechanisms by which viruses can become integrated into their hosts, akin to mobile elements, and as a result lead to persistence of those sequences with consequences for the host genome. In the case of HPV16, a small number of changes in its DNA may be responsible for differential host genomic instability and integration propensity, as evidenced by a difference in the rate of integration between HPV16 D2 and D3 sub-lineages [Lou *et al*., 2020]. Additional bioinformatics analysis of this study and other *ex vivo* data, coupled with *in vitro* experimental work, is required to further test the HPV integration hypothesis and the impact of other co-factors. Hypothesis-free exploration of these data will also be important to clarify the gaps in our existing knowledge and identify fascinating phenomena and molecular mechanisms to test in future experiments [Pipas, 2019]. From an applied science and clinical perspective, HPV16 sub-lineage genotyping can be used for preventative screening and routine monitoring, and next-generation sequencing specifically could be used on blood to detect integrated HPV sequences in circulating tumour DNA [Holmes *et al*., 2016] as a tumour-specific diagnostic biomarker and for monitoring residual disease and recurrence after treatments.

## METHODS

### Data access and acquisition

Access to controlled sample data from TCGA, using the database of Genotypes and Phenotypes (dbGaP), was acquired via application to the electronic Research Administration (eRA) commons. We registered our host institution, Lakehead University, as a new organization and received a Data Universal Numbering System (DUNS) identifier for access. Our proposal title was “Characterization of human papillomavirus type 16 integration sites in cervical and head and neck tumour biopsies”, with approval allowing access to TCGA Cervical Squamous Cell Carcinoma and Endocervical Adenocarcinoma (TCGA-CESC) as well as Head and Neck Squamous Cell Carcinoma (TCGA-HNSC) files for analysis. After receiving access to the data (a multi-step administrative process, outlined in **Figure 7A**) TCGA case files were transferred via their universally unique identifier (UUID) using the command-line implementation of the Genomics Data Commons Transfer Tool v1.4.0 and analyzed on the Compute Canada Cedar cluster, which uses the Secure Shell (SSH) encryption protocol for secure data transmission.

**Figure 7.**
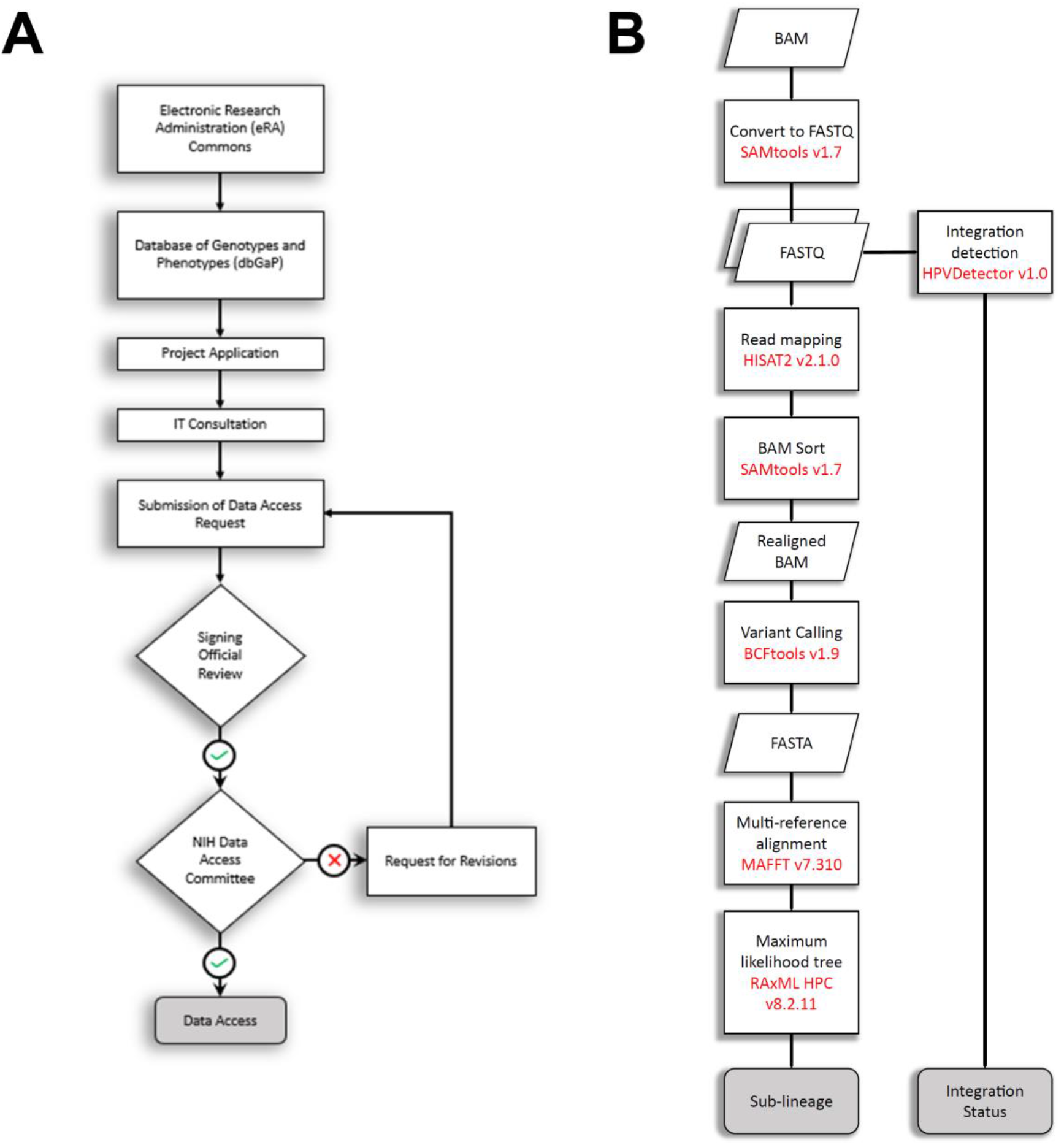
Access and analysis of TCGA data. **(A)** Access to controlled sample genomic and transcriptomic data from TCGA required a multi-step authorization process before being able to access relevant datasets. This administrative process, which requires annual renewal and project revision, takes approximately three months initially. **(B)** Our analytical workflow involved the following computational steps: TCGA high-throughput sequence read data (WXS or RNA-Seq) were first accessed and converted from pre-aligned BAM/SAM format to FASTQ read format. Data were then subjected to HPV16 sub-lineage determination and integration analysis.

High-throughput sequencing data from HPV16-positive cervical carcinoma-derived cell lines SiHa [Friedl *et al*., 1970] and CaSki [Pattillo *et al*., 1977] were used as controls. For SiHa, which contains low-copy number integrated HPV16, we used the same sample used in the HPVDetector paper [Chandrani *et al*., 2015], a whole-genome sequence from the DDBJ (DNA Data Bank of Japan) Sequence Read Archive (DRA) run SRR1609142, from study SRP048769. For CaSki, which contains a high-copy number of tandemly repeated HPV16 genome integrations, we downloaded RNA-Seq data from the Sequence Read Archive (SRA) run SRR9924750, from project PRJNA484129 [Sen *et al*., 2020].

### HPV16 positivity and sub-lineage genotyping analysis

While HPV16-positive cervical cancer (TCGA-CESC) samples had associated sub-lineages already identified (albeit, with 10 of the 16 current sub-lineage reference genomes [TCGA Research Network, 2017]), sub-lineage genotyping was not immediately available for the head and neck (TCGA-HNSC) cases [TCGA Research Network, 2015]. It was thus necessary to develop a pipeline for ascertaining the sub-lineages of unknown identified samples (*i*.*e*., those that were identified as HPV16 positive by Cantalupo *et al*., [2018] as well) in addition to confirming their status as HPV16 positive through an analysis of our own. Our pipeline follows an analytical workflow starting with the pre-aligned BAM files from TCGA [**Figure 7B**]. First, BAM files were converted into FASTQ files using SAMtools v1.7 [Li *et al*., 2009]. The reads were then mapped to the reference genomes of each HPV16 sub-lineage [**Table 2**] using HISAT2 v2.1.0 [Kim *et al*., 2015]. From the sixteen separate alignments performed, the one with the highest number of mapped reads was used for quantifying the presence of HPV16 by calculating the number of viral reads per million sequencing reads (RPM = 10^6^ x number of mapped reads/total reads, which accounts for sequence depth variation between sample libraries). RPM ranges for positivity analysis were based on a prior study which sought reasonable cutoffs for HPV detection in cancer tissues [Montgomery *et al*., 2016]. If RPM < 0.99, the sample was marked as negative across all categories and the analysis was terminated. Samples with RPM between 0.99 and 99 were marked as having “low” positivity for HPV16, while samples of RPM > 99 were marked as having “high” positivity. Given their large range, all RPM values were subjected to an inverse hyperbolic sine (IHS) transformation [Burbidge *et al*, 1988] following IHS(x) = log_10_(x + (x^2^ + 1)^1/2^), an alternative to traditional logarithmic transformation that allows values lower than 1 to have a plottable log value and which has seen recent use when analyzing biological data [Haines *et al*., 2020]. SAMtools v1.7 was then used to sort and prepare the files corresponding to these samples for conversion to VCF format. BCFtools v1.9 [Li, 2011] was used for variant calling and generating the consensus sequence. The new consensus sequences along with all reference sequences were aligned against each other using MAFFT v7.310 [Katoh *et al*., 2009] with default parameters. Maximum likelihood trees were inferred using RAxML HPC v8.2.11 [Stamatakis, 2014], under the general time-reversible nucleotide model with gamma-distributed rate heterogeneity and invariant sites (GTRGAMMAI). Sub-lineages were determined from Newick tree files using ETE Toolkit v3.0.0b34 [Huerta-Cepas *et al*., 2010] based on topological distance. Each sample was incorporated into the reference tree [**Figure 8**] and assigned a sub-lineage corresponding to the nearest neighbour.

**Table 2.** HPV16 sub-lineage reference genomes used for genotyping. Derived from reference data summarized in Burk *et al*. [2013], Chen *et al*. [2018], and Mirabello *et al*. [2018].

| HPV16 sub-lineage | GenBank # | Other ID | Original nomenclature |
| --- | --- | --- | --- |
| A1 | K02718 | NC_001526.4 (latest) | European Prototype (EP) |
| A2 | AF536179 | w0122 | European (E) |
| A3 | HQ644236 | AS411 | European (E) |
| A4 | AF534061 | w0724 | Asian (As) |
| B1 | AF536180 | w0236 | African-1a (Af-1a) |
| B2 | KU053915 | IARC1100085NI | African-1 (Af-1) |
| B3 | HQ644298 | Z109 | African-1b (Af-1b) |
| B4 | KU053914 | IARC907912AL | African-1 (Af-1) |
| C1 | AF472509 | R460 | African-2 (Af-2) |
| C2 | HQ644244 | IARC240211GU | African-2 (Af-2) |
| C3 | KU053920 | IARC040105BE | African-2 (Af-2) |
| C4 | KU053925 | IARC304612MO | African-2 (Af-2) |
| D1 | HQ644257 | Qv00512 | North-American-1 (NA1) |
| D2 | AY686579 | Qv15321 | Asian-American (AA) 2 |
| D3 | AF402678 | Qv00995 | Asian-American (AA) 1 |
| D4 | KU053931 | IARC903812AL | North-American-2 (NA2) |

**Figure 8.**
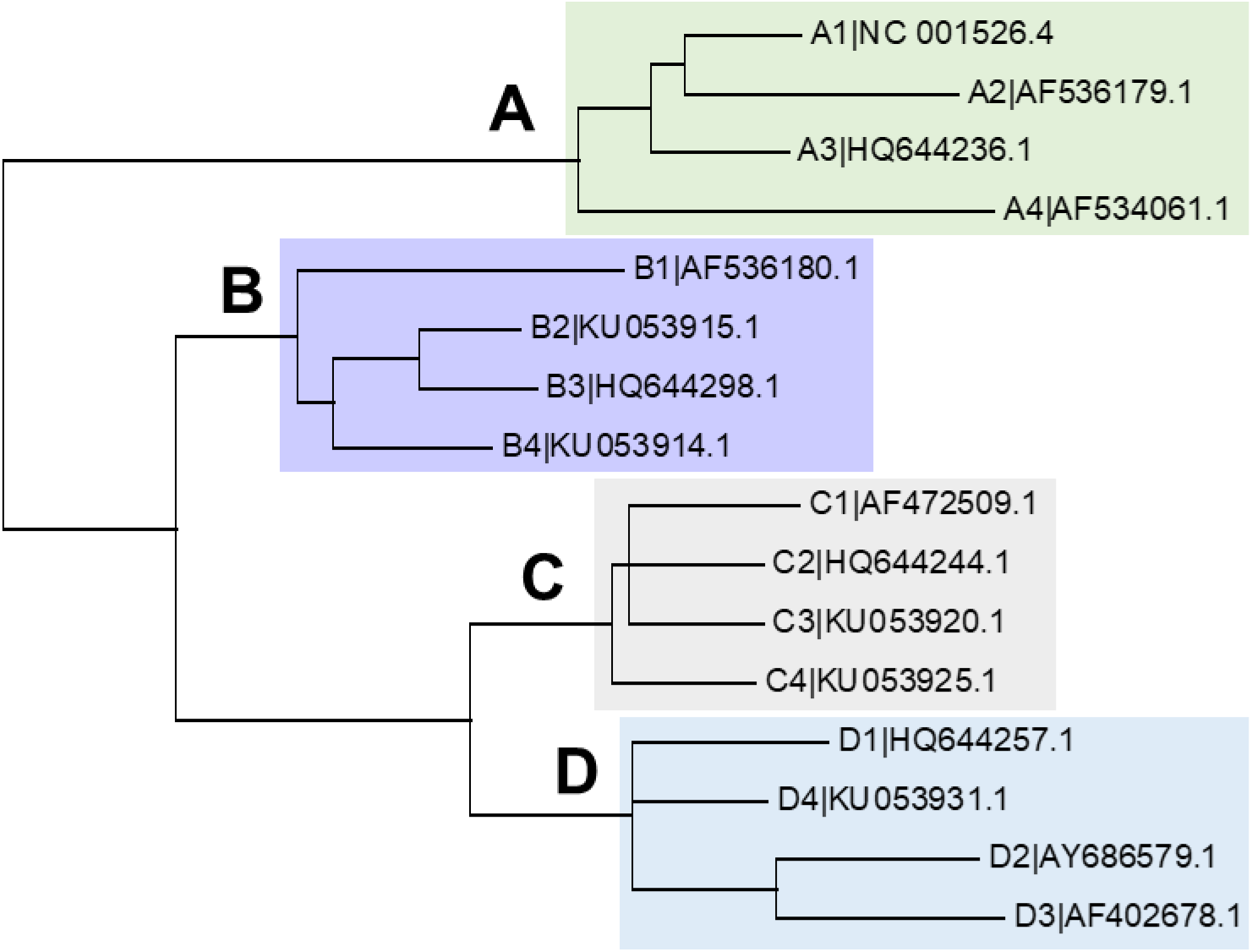
HPV16 sub-lineage reference tree for phylogenetic analyses. HPV16 phylogenetic maximum likelihood tree used as a reference for sub-lineage determination. Branches correspond to four lineages (A, B, C, and D), each containing four sub-lineage genomes (1 to 4). The tree was built with FigTree v1.4.4 using standard settings.

### Viral-human integration detection and characterization

Several open-source software tools for integration analysis are available for WGS, WXS, and whole-transcriptome sequencing (RNA-Seq): *e*.*g*., VirusSeq [Chen *et al*., 2013], ViralFusionSeq [Li *et al*., 2013], VirusFinder [Wang *et al*., 2013], HPVDetector v1.0 [Chandrani *et al*., 2015], ViFi [Nguyen *et al*., 2018] and VirTect [Xia *et al*., 2019]. The latter tool name, VirTect, was also used by another group for viral genome sequencing and host genes related to carcinogenesis using RNA-Seq [Khan *et al*., 2018]. HPVDetector v1.0 was chosen since it was specifically designed for detecting and annotating HPV integration events, as well its compatibility with the Compute Canada Cedar cluster environment. VirTect was also considered but is currently incompatible with available modules in Cedar, requiring an older version of Python (Python 2.7 vs 3.0). Integration analysis was performed using command-line execution of HPVDetector v1.0 [**Figure 7B**] set to “Integration” mode for all samples and “Transcriptome” or “Exome” mode depending if the sample contained WXS or RNA-Seq data, respectively. Raw output was obtained in the form of the number of integrated reads. This value was then converted to a percentage of the total number of viral reads to account for sequencing depth variation amongst selected samples and used as a metric to quantify viral-host integration. Samples with values at or above 1.14% were classified as positive for HPV16 integration, a threshold obtained through comparison to the landmark CESC study cases to maximize concordance. The remaining samples were considered as positive for the episomal form of HPV16.

## DECLARATIONS

## Acknowledgements

The results reported here are based upon data generated by The Cancer Genome Atlas Research Network (cancergenome.nih.gov) and we are grateful to the participants and Network members for this resource. This research was enabled in part by computational support provided by Compute Ontario (www.computeontario.ca) and Compute Canada (www.computecanada.ca). Thank you to Darryl Willick and Dr. Wely Floriano for their assistance in helping set up and maintain required bioinformatics tools (such as the Galaxy platform) hosted at the Lakehead University High Performance Computing Centre (LUHPCC) as well as Dr. Zigui Chen who shared a HPV sub-lineage detection pipeline. We are also thankful for Bruce Bogacki’s technical support at the Thunder Bay Regional Health Research Institute (TBRHRI). Additional thanks to Kathleen Roulston, Vanessa Masters, Anirudh Shahi, Dallas Nygard, and Christopher Gibb for assisting with preliminary literature review, initial data access, and early analyses. Preliminary work related to this study was presented as a poster in Heidelberg, Germany (11-14 Oct 2017) at The Mobile Genome: Genetic and Physiological Impacts of Transposable Elements conference and talks in Thunder Bay, Canada at the Health & Information Technology (HIT) Research Group Workshop (12 Oct 2018) and the Health & Information Technology & Biotechnology & Allied Sciences (HITBASS) Symposium (27 Oct 2019). This work was supported by the Natural Sciences and Engineering Research Council of Canada (NSERC) grants to IZ (#RGPIN-2015-03855) and RJ (CGS-D#454402-2014). The funding body had no role in study design, data collection, data analysis and interpretation, or preparation of the manuscript. We declare no conflicts of interest.

## Authors’ contributions

The study was initially conceived of and designed by IZ and RJ. Data access was managed by RJ, IZ, and AOV. Methods were developed, optimized, and analyses performed by AOV with input from RJ and IZ. Biological interpretation, literature searches and manuscript writing were led by RJ and IZ, with input from AOV, while all authors were involved in providing critical revision and feedback.

## Notes

### Competing Interest Statement

The authors have declared no competing interest.

### Summary of Updates

Additional cases/samples included; qualitative analyses expanded.

